# iCAVE: an open source tool for immersive 3D visualization of complex biomolecular interaction networks

**DOI:** 10.1101/061374

**Authors:** Vaja Liluashvili, Selim Kalayci, Eugene Flouder, Manda Wilson, Aaron Gabow, Zeynep H. Gümü

## Abstract

Visualizations of biomolecular networks assist in systems-level data exploration in myriad cellular processes in health and disease. While these networks are increasingly informed by data generated from high-throughout (HT) experiments, current tools do not adequately scale with concomitant increase in their size and complexity. We present an open-source software platform, interactome-CAVE, (iCAVE), that leverages stereoscopic (3D) immersive display technologies for visualizing complex biomolecular interaction networks. Users can explore networks (i) in 3D in any computer and (ii) in *immersive* 3D in any computer with an appropriate graphics card as well as in CAVE environments. iCAVE includes new 3D network layout algorithms in addition to extensions of known 2D network layout, clustering and edge-bundling algorithms to the 3D space, to assist in understanding the underlying structures in large, dense, layered or clustered networks. Users can perform simultaneous queries of several databases within iCAVE or visualize their own networks (e.g. disease, drug, protein, metabolite, phenotype, genotype) utilizing directionality, weight or other properties by using different property settings. iCAVE has modular structure to allow rapid development by the addition of algorithms, datasets or features without affecting other parts of the code. Overall, iCAVE is a freely available open source tool to help gain novel insights from complex HT datasets.

## Introduction

Interaction networks are one of the primary visual metaphors for communicating and understanding -omics data at a systems level. From cellular organisms to human society, they provide critical clues on systems-level behavior^1–3^, and in biomedicine they are essential for understanding both normal^4,5^ and disease states^6–9^, and instrumental for drug discovery^10–12^ as well as biomarker identification^13–15^. Changes in networks have been shown to help in prognosis for breast cancer patients^6^, analyzing systematic inflammation in humans^8^, or studying emerging tumor markers^16^. Network visualizations have thus become key tools in basic and translational biomedical research. Consequently, there is an abundance of tools for their interpretation and exploration^17,18^. Many of these tools are also coupled with public databases, allowing data visualization and interpretation in the context of previous knowledge^17^. In fact, currently there are more than five hundred resources listed at http://pathguide.org that provide access to thousands of networks, cataloging millions of interactions between biomolecules^19^.

Among the currently available biomolecular network visualization tools, the most popular is an open-source and freely available tool, Cytoscape^20^, which enables explorations with different filter, layout, color and cluster options, and includes estimations of network topology parameters and centrality measures. There are also a number of JavaScript network visualization libraries (e.g. sigma.js http://sigmajs.org/), and software packages (e.g. iGraph http://igraph.org) on the web. However, the currently available layout algorithms in these libraries and employed in Cytoscape^21^, in addition to other tools like Ingenuity^22^, Osprey^23^, VisANT^24^, BINA^25^ to name a few, are inherently limited by the number of molecules and interactions that can be displayed in the 2D-space of a screen, and the associated layout and representation challenges. As the databases that supplement biomolecular interaction networks are growing at an unprecedented rate due to the increasing size and complexity of -omics experimental techniques, innovations are necessary to address the challenges the concomitant large, dense, and/or multi-layered networks present. Furthermore, because increasingly powerful technologies have enabled the collection of data from multiple types of cellular events simultaneously, in order to achieve better understandings of such complex processes, it may be necessary to maximally integrate data across multiple dimensions, pushing the limits of current visualization tools.

New visualization solutions can bring substantial benefits by improving our understanding of complex mechanisms in human disease, reducing the time to discovery and diagnosis. One approach has been to shift to 3D, and even to immersive 3D space by employing cave automatic VR environments (CAVEs), which are immersive VR environments that include projectors directed to several walls of a room-sized cube^26^. While the relative benefits of immersive 3D in network visualization are still being debated^27–29^, a recent study that visualized the same network in an immersive 3D CAVE environment *vs*. 2D display identified a global network property due to the additional features of the CAVE, and quantitatively validated this result by comparing to 1000 random permutations of networks of the same size and distribution^26^. These beneficial features were stereoscopic visualization, magnification and wide field-of-view^26^. Stereoscopic visualization creates the illusion that objects seen are volumes in 3D-space, which results from the projection of separate left and right eye images of each object, and then combining them in stereo-enabled eyeglasses. Additionally, motion sensors on the eyeglasses enable automatic detection of the user’s location if she is moving, and adjust the image perspective, and hand-held controls enable user-network interactions such as zooming and rotating the view. However, the study has not introduced a tool to the community. Also, the researchers only utilized an extension of the standard force-directed network layout of Fruchterman-Reingold^30^ to 3D, and did not test performance of additional existing or new layout or clustering algorithms or topology parameter measurements. To perform similar comparative immersive 3D *vs*. 2D studies, to test alternative algorithms or to analyze networks in immersive 3D other researchers would need to write their own code. They would also need a CAVE, which is a substantial investment to build and maintain.

Note that while several network visualization tools incorporate 3D layouts^31–33^, they are **not immersive 3D**, meaning that they do not have interoperation capability with Virtual Reality (VR) technologies, and their displays are in 2D. For example, Arena 3D^31^ mixes 3D properties with 2D, by arranging data in multilayered graphs in 2D, with each layer representing a different data type. While the tool then implements several layout and clustering algorithms for each layer, and layers can be zoomed in/out and rotated, it does not offer global layout and clustering algorithms to make full use of the third dimension: each layer still has a 2D layout on its surface. Directed edges are also not supported^31^. 3DScapeCS^32^ is a Cytoscape PlugIn written in Java, with built-in network layout algorithms that are extensions of the classic 2D force-directed layouts. The tool does not allow users to add new layouts or functionalities^30^ and does not utilize 3D effects to help improve comprehension, such as transparency, or advanced shadow effects. BioLayoutExpress^33^ (current name Miru) is a stand-alone 3D application specifically for gene expression networks, which currently provides three network layout algorithms, a single clustering method, no edge bundling and a limited number of network topology statistics, which cannot be saved by the user and does not allow directional edges. Importantly, the tool has a licensing fee and thus is not freely available.

In summary, 3D network visualization field is still somewhat nascent. We need freely available open-source tools for biologists to visualize their biomolecular networks, and at the same time for algorithm developers to add and test their methods that take advantage of the third dimension. This will help the community to understand how best to exploit features unique to 3D in biomolecular network exploration, providing insights to the ongoing debate on the advantages of (immersive) 3D *vs*. 2D.

Here, we introduce interactome-CAVE (iCAVE), an open-source tool for 3D and immersive 3D visualization of complex networks. It is designed primarily to assist biomedical researchers in data exploration, though it can be used in any field that involves networks. iCAVE development is made possible by the continuous evolution of data analysis tools in VR, stereoscopic visualization and emerging 3D technologies. Use of VR technology in life sciences research is still nascent^34–37^, and so far do not include freely available open source tools for biomolecular network visualizations, mainly due to the limited portability of the technology to personal computers until recently. We designed iCAVE without this limitation, by taking advantage of recent advances in computer graphics hardware, software and content creation that are leading to a proliferation of stereoscopic visualization capabilities in personal computing. Driven primarily by gaming and movie industries, computers can now be upgraded to display high quality stereoscopic 3D visuals with wireless glasses and advanced software^38^ at nominal prices. As most scientific computers are becoming 3D-capable and the glasses are going mainstream, iCAVE is on the leading edge of this larger trend in the evolution of visual computing technology. While iCAVE works in CAVE environments, its real benefit comes from enabling immersive 3D network visualization in stereo-enabled computers. Therefore, if a computer is equipped with stereo capabilities, users can display immersive 3D visualizations. If the computer is not equipped with stereo capabilities (or if users choose to turn off stereo), iCAVE provides an interactive (non-immersive) 3D environment that still offers most of its features.

As a visualization and analysis tool, iCAVE enables network explorations in hypothesis-driven contexts that is flexible, collaborative and user friendly. It introduces new 3D algorithms that are built-in for laying out nodes and their connections in 3D space (hemispherical and multi-level layouts) as well as graph clustering algorithms for clustering the nodes based on network structure and connectivity and then laying out the resulting clusters in 3D space. It also allows the users to add their own layout or clustering algorithms. Furthermore, users can visually integrate multiple clusters or data types from several databases within the same graph as a multi-layered network (e.g. metabolomic, proteomic, genomic, GWAS-disease, protein-drug interactions). iCAVE reports several network topological properties and centrality measure statistics as 2D reports (now shown). While not extensive, it also includes several built-in databases, to assist in preliminary mapping of High-Throughput (HT) experimental data in the early discovery phase of network building. Customizable color, texture, size and layout options assist in displaying maximum information in a graph in an optimized manner. Edges can be in user-defined colors, weights and directions and can be bundled together for simplified views. Data can be uploaded to iCAVE in a simple tab-delimited text file format; output can be saved as 2D snapshots or movies configured with user-defined rotation, zoom and speed parameters.

## Results

ICAVE users can utilize features that are unique to 3D or immersive 3D visualizations and test whether these improve the quality of their network exploration. These features include stereoscopic visualization, wide field of view, magnification, motion sensors and hand-held controls, as described in introduction section. For example, consider rendering a 2D biomolecular network affected by genomic alterations in glioblastoma^39^. In this example, the network layout algorithm is a simple 3D extension of the classical force-directed Fruchterman-Reingold^30^ (Fig 1). Note that instead of the static 2D network in Fig. 1A, iCAVE users experience full 3D depth perception at the comfort of their own stereo-equipped computer (Fig. 1B), or inside a CAVE (Fig. 1D). Furthermore, users without a stereo-equipped computer are also able interact with the 3D network: by using their mouse (in lieu of hand-held controls), they can zoom or rotate the network to a view without occlusions. This ability to rotate and zoom enables viewing of the network from different view angles, such as the screenshot in Fig. 1C. Such easy exploration enables users to visually identify a feature unique to the topology of this example network. The network feature was not intuitive from the original 2D layout in Fig. 1A: nodes CBL and SPRY2 (with *) are *connectors* between two dense network regions (modules) (Fig 1A-C). A targeted attack to these genes can split the network into two. Such discoveries of network topological features, among others, give a richer, more intuitive and ultimately more insightful understanding of network data.

**Figure 1.**
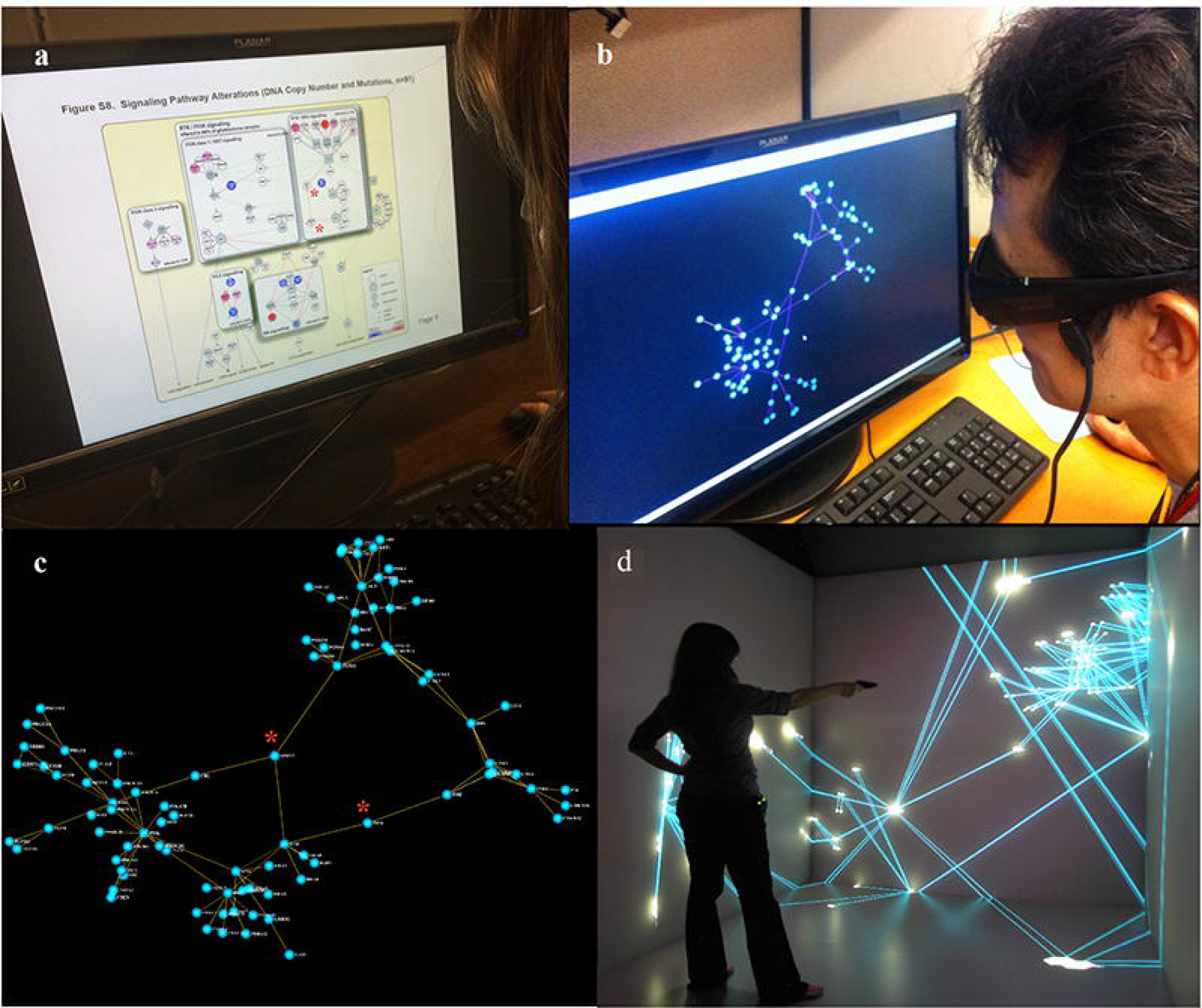
Comparison of various displays. **a.** User interacting with a flat 2D display of manually curated pathways most affected by genomic alterations in glioblastoma ^71^. **b.** User experiencing a full 3D depth perception of the same network with iCAVE while using stereoscopic glasses on his desktop. iCAVE display is generated with force-directed layout algorithm. Notions of edge crossings that create a hairball effect in 2D have little meaning in 3D, as the user can navigate to a view without occlusions, solving the visual clutter problem. **c.** A screenshot from the 3D display generated with the force-directed layout. This network is generated without *a priori* knowledge of the network topology; however, it readily identifies hubs, connectors and modules, such as the connectors between two dense regions of the network (highlighted with a (*) in both panels **a** and **c). d.** Immersive visualization in a CAVE environment, with one user inside the data space. While the photos only capture images reflected on the interaction walls of the CAVE, the user is interacting with a virtual 3D image. In both **b** and **d,** the options to zoom and rotate the network with a mouse (or wand) click helps the user focus on a particular hub or module of interest within the network. While the addition of the third dimension gives a richer, more intuitive and ultimately more meaningful understanding of the network-represented data, the 3D layout brings a completely new modality into network visualizations, with clean, easy to use and understandable layouts.

### Addressing Large Networks

When exploring large networks, researchers may miss important characteristics if they cannot interact with the complete network. In the simplest case, the nodes may form (i) dense sub-networks that are interconnected by a small number of *connector* nodes which render them critical or (ii) multiple networks (often one giant and few smaller ones) where the smaller sub-networks may represent functional groups of importance, such as a critical enzyme complexes. Thus, there are benefits visualizing the complete network even if it is very large. At the same time, while the human brain has a remarkable capacity to visually identify patterns, enabling interpretation of data, visualizations of large networks may exhibit problems with display clutter, molecular positioning or perceptual tension, which may lead the user to misinterpret closely positioned molecules as related^47^. Such misinterpretations are inherent in the limitations of human visual perception, and have been well-studied in (Gestalt) psychology: people tend to organize visual elements into groups^48^.

Using 3D layouts, the elements that appear to form a pattern because of their visual positioning in one viewpoint can be interpreted correctly by rotating the image to a different viewpoint (as shown in Fig. 1). Furthermore, in networks that are denser or larger than that of Fig.1, the potential 2D *hairball* effect can obscure important interactions. Using iCAVE, the user can simply navigate to a view without occlusions by moving her head, rotating the image, and zooming in or out, so that *edge-crossings* causing the *hairball effect* in 2D are eliminated. To further address cluttering problem, iCAVE provides an *edge-bundled display*^49^ option for visually bundling adjacent edges together, analogous to bundling of electrical wires or cables. Bundling is extremely useful in identifying global patterns in very large networks and can suggest vulnerabilities as targets. There are several layout algorithms built-in within iCAVE to address the molecular positioning problem; depending on the topology of the network to be visualized, one layout may work better than another. We suggest testing each layout to see which works best. We provide examples of how these features can help with exploring a network in the following sections.

***New biological insights gained from networks with known 3D physical coordinates*.** Users can generate visualizations of physically constrained networks at multiple scales, ranging from proteins (Fig. 2A) to the whole brain (Fig. 2B). Visualizations that employ physical positions of a 3D network coupled with edge bundling can provide insights during hypothesis generation.

**Figure 2.**
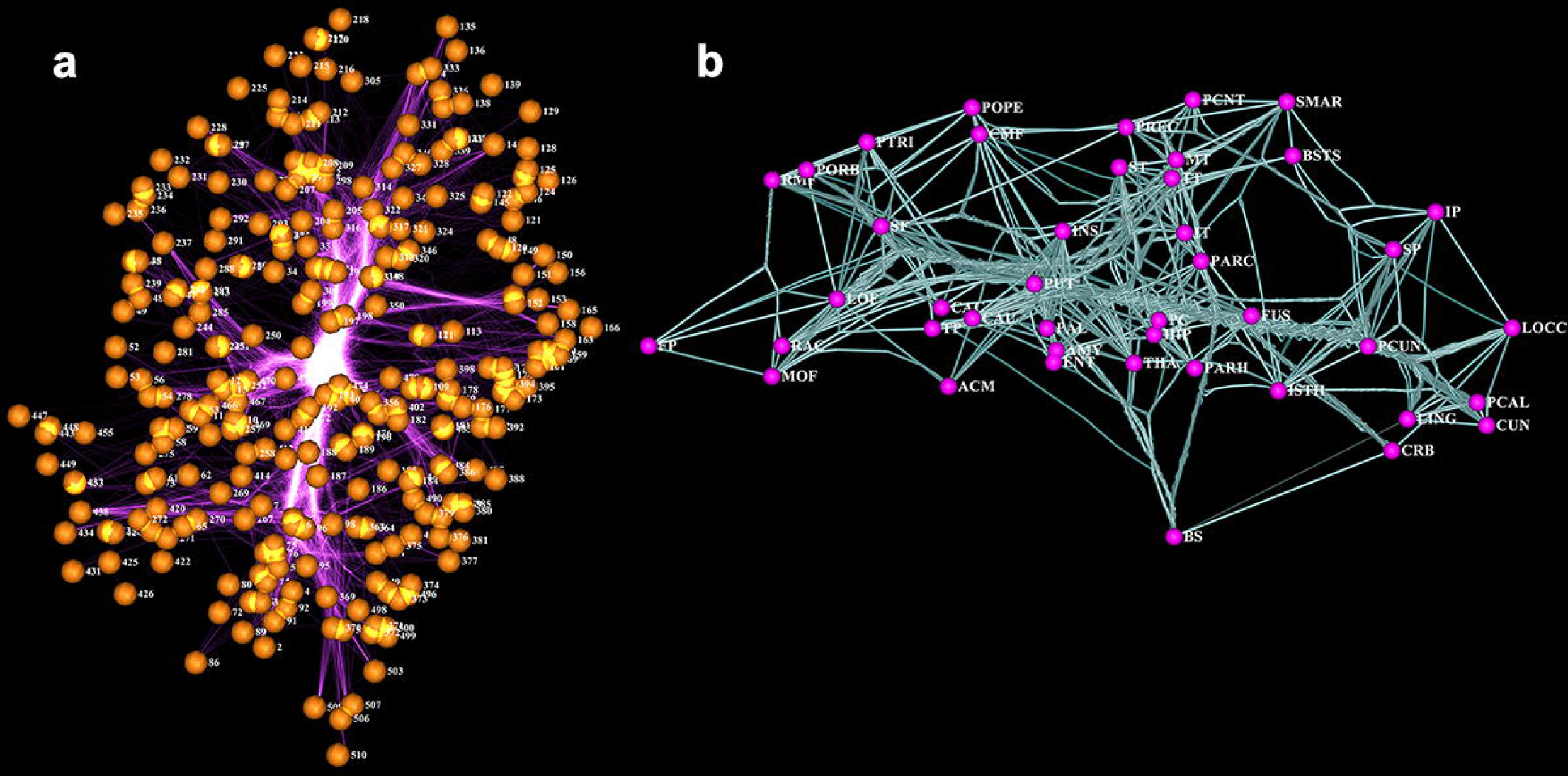
a. iCAVE visualization of bacterial leucine transporter, LeuT residue correlation network, side-view. Nodes represent 3D coordinates of alpha-carbon of a residue; edges represent top 3,000 (Pearson) correlations between residue pairs, where the input is 3D coordinates & correlation scores. Surprisingly, 3D visualization with edge bundling enables representation of highest density correlations (*correlation highways*) that travel through the substrate permeation pore in protein center, connecting extracellular and intracellular domains. Correlation highways at the pore are visually fascinating and biophysically intuitive. Information highways of some residues outside the pore reveal unexpected structural importance (data courtesy of Weinstein Lab, Weill-Cornell). **b.** Living Human Brain Connectivity. iCAVE visualization of brain regions as nodes, labeled by anatomical region name. Edges show connectivity, and bundling shows *connectivity highways*. Datasets from Diffusion Tensor Imaging of left hemisphere scanned ith Siemens 1.5Tesla and generated by Fiber Assignment by Continuous Tracking tractography using U. of California, Los Angeles (UCLA) Multimodal Connectivity Package connectivity matrix module. Database is powered by the Human Connectome Project, which aims to provide an unparalleled compilation of neural data to achieve never before realized conclusions on the living human brain.

For example, Fig. 2A represents a snapshot of bacterial leucine transporter (LeuT) residue correlation network, where the nodes represent 3D coordinates of alpha-carbon of a residue and edges represent top 3,000 (Pearson) correlations between residue pairs from a Molecular Dynamics simulation (from Michael LeVine, personal communication). Remarkably, bundling the edges of this network enables the representation of highest density *correlation highways* that travel through substrate permeation core in protein center, connecting extracellular and intracellular domains. While these highways are visually fascinating, they also enable users to identify specific residues that have dense correlation highways outside of the protein core, which are unexpected. These residues may have previously unidentified importance in protein structure and are potential candidates for follow-up studies.

### Automated Layouts utilize 3D for molecular positioning

The topology of cellular and disease networks tend to follow basic and reproducible organizing principles, and navigating the entire network provides a good initial understanding of such a network. The network layout algorithm must address the complex problem of arranging the nodes to clearly disseminate the topology, and at the same time be visually pleasant and user-friendly. iCAVE offers several algorithmic options for network layout to achieve these aims.

Due to user familiarity, we extended several variations of the force-directed layout approach to 3D: (i) the classical *force-directed algorithm*^30^ treats the network as a physical system with edges analogous to *springs* and nodes to *electrically charged particles* that repel each other. The final layout is established at the state at which the repulsive and attractive forces balance each other^40^ (see Figure 3); (ii) *Lin-log layout*^41^ is better suited for larger networks because it keeps highly connected nodes in close proximity with minimal number of edge crossings; and (iii) *hybrid force-directed layout*^42^ partitions the graph into smaller units before applying the force-directed algorithm (see Methods). We further implemented two novel layout algorithms to take full advantage of immersive 3D:

**Figure 3.**
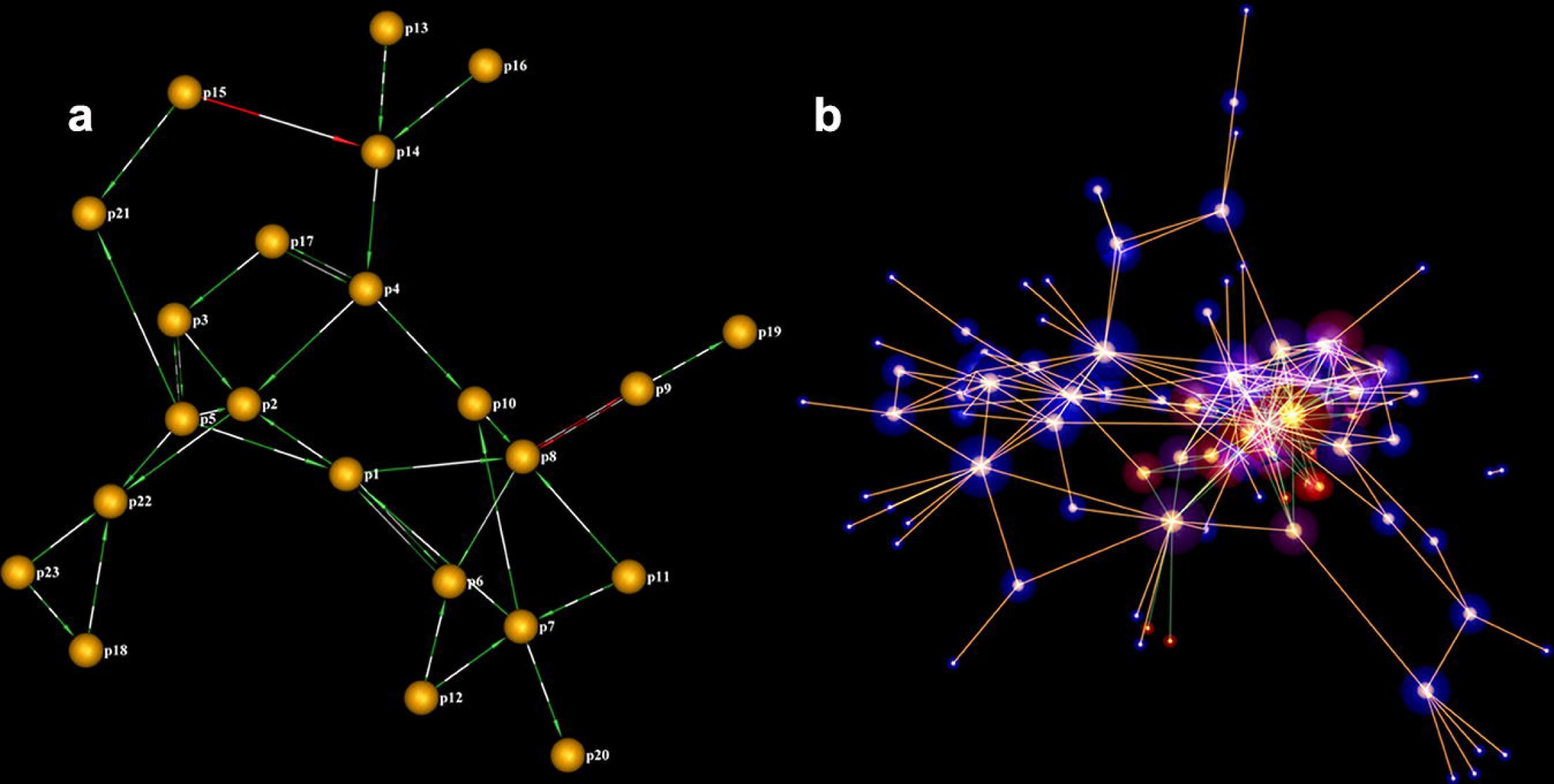
a. A directed and weighted signaling network using semantic layout. Edge color represents activation (red) or repression (green). Color alteration frequency represents weight (Data courtesy of Chris Sander, Memorial Sloan Kettering). **b. 3D iCAVE force-directed layout representation of a bacterial network.** Nodes are bacterial species, edges are top nonlinear between-species relationships. Node size is proportional to the number of relationships per node. Orange edge color: relationship explained by diet, node glow: fraction of orange edges (100% is red; 0% is blue. Data: ^72^).

*Semantic levels layout algorithm* segregates the network into separate layers (default 7) in the third dimension. The layout within individual layer is calculated using a 3D extension of the force-directed approach. Semantic layers layout can be especially useful for user-defined networks where the number of layers and node assignments to layers can correspond to different data types (e.g. see a 2D projection in Figure 5 and 3D video in Supplementary Video 3, where layer1: genes; layer2: diseases; layer3: drugs).

*Hemispherical layout* is a novel layout algorithm we have developed, that positions the network on the surface of a 3D hemisphere. The most connected node is positioned at the top center of the hemisphere. Then, the whole hemisphere surface is populated based on a decreasing rank-order of connectivity. The node positions are fixed and the edges are drawn on the hemisphere surface (e.g. see a 2D projection in Figure 6C and 3D video in Supplementary Video 4).

Each layout algorithm has unique strengths and we recommend the user to test different options. Semantic layout is often ideal for hierarchical networks. Force-directed layout often captures the essence of large networks. Hemispherical layout leads to clean images with optional edge bundling (Figure 6C and Supplementary Video 4).

### Statistics on Network Topological Properties

Most real-world networks exhibit substantial and non-trivial topological features, where connections are neither purely regular nor random. iCAVE automatically generates and reports network topology statistics and centrality measures both graphically and in tabular form (not shown). These statistics include the number of nodes, the number of edges, network diameter, node-betweenness centrality, closeness centrality, neighborhood connectivity, shortest path, topological coefficient, and node degree distribution properties of the network.

### COMBO Database for Simultaneous Query of Multiple Data Types

The currently publicly available biomolecular interaction data are often contained in databases that are massive in size^19^. While not comprehensive, iCAVE combines data from multiple resources into a single COMBO repository to enable quick queries. This includes protein-protein interaction databases Human Protein Reference Database (http://www.hprd.org) and intAct (http://www.ebi.ac.uk/intact), disease and associated gene variants database (http://www.genome.gov/gwastudies); and drug-target databases STITCH (http://stich.embl.de) and DRUGBANK (http://www.drugbank.ca). Pathways database SuperPathway is stored separately (personal communication with Josh Stuart, UCSD). Users can add their own databases without affecting other parts of the code. Details on the COMBO database are given in Supplementary Table 1.

### Visualizing Multiple Layers of Information

Effective usage of genomic information can depend on finding systems-level connections between multiple types of information, such as that of between genomic variation, disease and drugs^43–46^. Visualizing such data by using semantic layout can assist in exploration in higher-level organization, all in one graph. User can pick a gene (e.g. AHR, dark blue, Fig 5A), query the COMBO database for diseases associated with its variants (purple); identify drugs that target it (green) and drug candidates that may target (light blue) due to guilt by association for having common targets with AHR-targeting drugs. These serve as initial candidates for subsequent binding site characterization. Querying COMBO database further generates a hierarchical network of proteins that interact with AHR (Figure 5B, middle layer), diseases associated with gene variants of AHR-interacting proteins (purple) and AHR targeting drugs (green).

### Illustrative Examples

**Example 1.** Visualizing the complete global network, even if it is very large, can enable visual identification of a pattern. For example, consider a large probabilistic causal network constructed from human omental adipose tissue in a morbidly obese patient cohort in Fig. 4A. The network consists of 7,601 nodes, 13,979 edges^50^. Nodes are the genes expressed in tissue; edges are derived from a Bayesian network reconstruction algorithm that leverages DNA variation for causality. Here, we highlight nodes that represent a signature of genes causally associated with inflammatory bowel disease (IBD) SNPs or disease pathways. Notice that within this global view of the massive network, there is a pattern of the IBD genes clustering together, which visually supports the hypothesis of functional relatedness.

**Figure 4.**
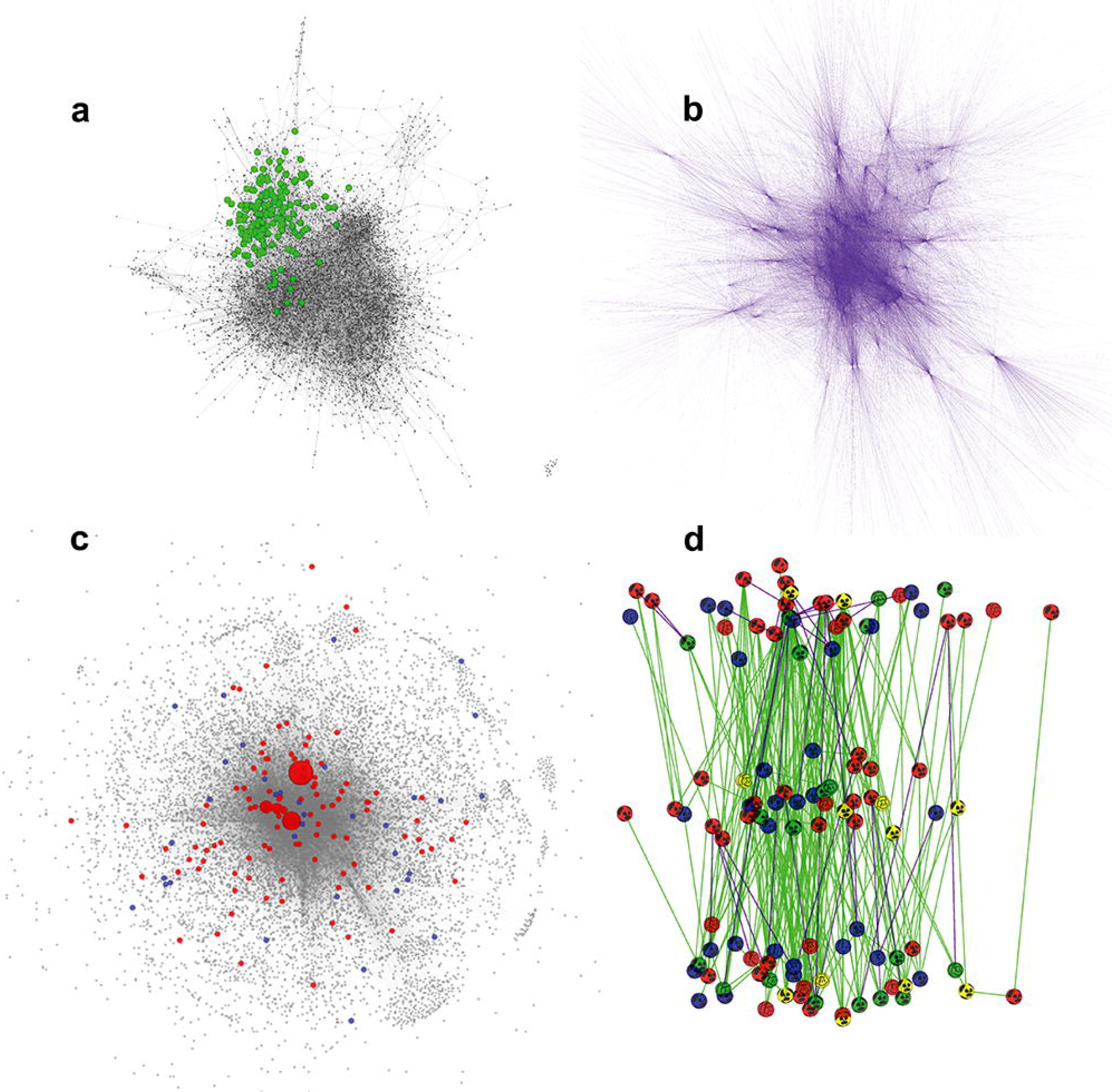
iCAVE print-ready images of massive networks in 2D with white background. **a.** Large probabilistic causal network constructed from human omental adipose tissue in a morbidly obese patient cohort (7,601 nodes, 13,979 edges)^50^. Nodes are gene expression traits in tissue; edges are derived from a Bayesian network reconstruction algorithm that leverages DNA variation for causality. Highlighted nodes represent a gene signature causally associated with disease SNPs or pathways. Signature genes clearly cluster together, suggesting functional relatedness. **b.** A network of 119 transcription factors (TFs), their 26,037 target interactions (edges) with 9,057 genes (nodes)^51^ from ENCODE study. **c.** A massive unified 'Multinet' of PPI, phosphorylation, metabolic, signaling, genetic and regulatory networks (14,558 nodes, 109,597 edges). Multinet correlates tolerance to loss-of-function (LoF) mutations and evolutionary conservation, with nodes for (LoF) tolerant (blue) and essential genes (red) easily distinguishable. Node size is based on the degree centrality of a gene. While essential genes tend to be bigger and central, LoF-tolerant genes are smaller in the periphery. **d.** A hierarchical network integrates TF, ncRNA, miRNA and protein-protein interaction (PPI) information. Hierarchy levels are based on the mutual relationships between TFs. Connectivity and hierarchy reflects genomic properties (top level TF-binding correlates with target expression; mid-level contains information flow bottlenecks and connections with miRNA and distal regions, revealing ideal drug targets) (data from: Marc Gerstein, personal communication). While the original 2D figure cannot display the interconnections between elements within the same hierarchical level, it is straightforward with the iCAVE semantic levels layout.

**Figure 5.**
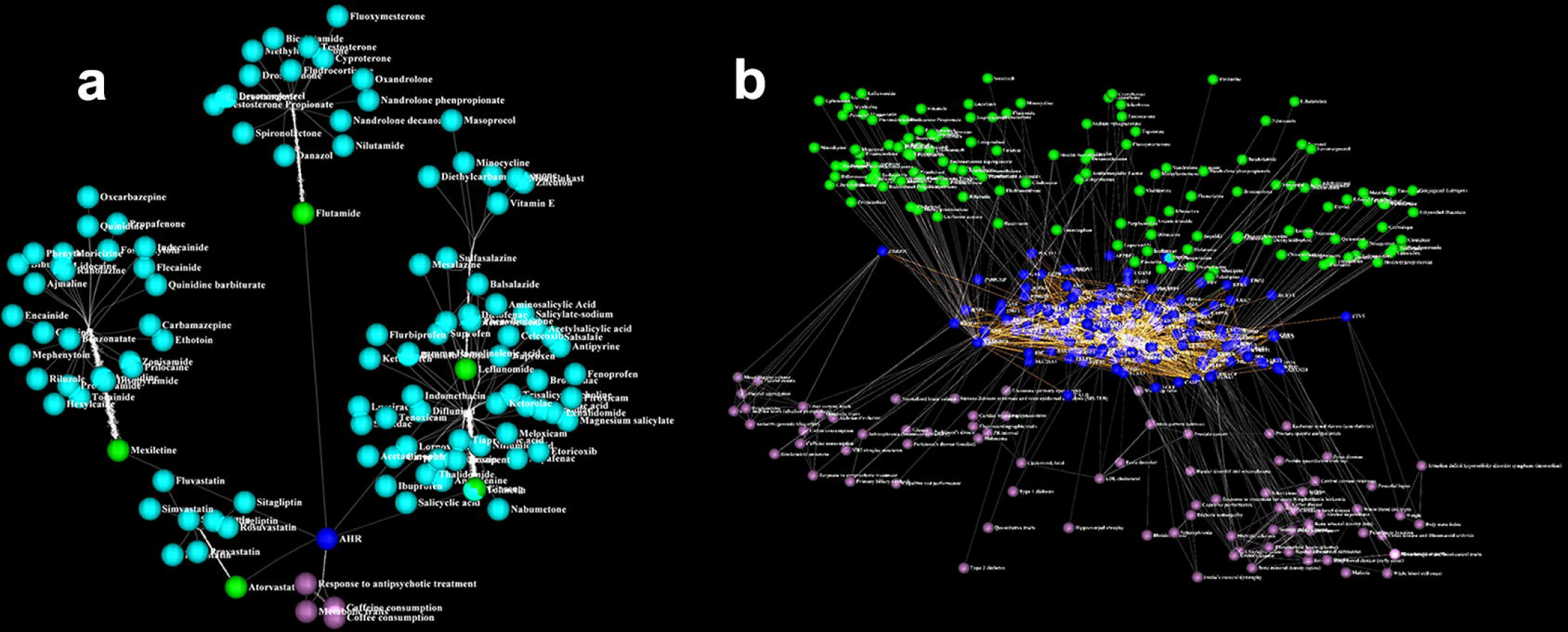
Visualizing Multiple Layers of Information. **a.** Using the iCAVE interface, we can pick a gene of interest (e.g. AHR, dark blue), and query the COMBO database for diseases that have been associated with AHR variants from GWAS studies (purple); drugs that are known to directly target AHR (green); and drug candidates that may directly interact with AHR (light blue). These drugs serve as an initial screening list of candidates for subsequent AHR binding site characterization. Semantics Levels layout segregates the layers. **b.** We can further query the COMBO database to generate a protein-protein interaction map of AHR (dark blue nodes; middle layer) and visualize the diseases associated with known SNPs (Single Nucleotide Polymorphisms) in the genes that code for AHR-interacting proteins (purple) and the drugs that directly target them (green). We provide a more detailed movie of this three-level semantics network with legible disease, gene and drug names in the Supplementary Video 3. In both Panels a and b, the user can click on any edge or node for further information (e.g. exact variant location for a specific disease from GWAS studies).

**Example 2.** While force-directed layout algorithms can help identify global patterns as in previous example, if the interaction network has a hierarchy, the semantic layers layout can help visualize the hierarchical nature of the interactions easily. For example, Fig 4B displays the global view of network generated from The Encyclopedia of DNA Elements (ENCODE https://www.encodeproject.org/) study data. The ENCODE Consortium is generating a comprehensive parts list of the human genome functional elements, including those that control active genes, such as transcription factors (TFs). Utilizing these unprecedented volumes of data, Gerstein and co-workers have generated the massive network in Fig 4B that includes 119 TFs that target 9,057 genes (nodes) via 26,037 interactions (edges)^51^. Using force-directed layouts, users can capture the general network structure and differentiate a TF from its neighbors by zooming in/out, adding labels to that specific TF, etc, as well as obtain statistics on its network centrality and other global topological properties as they pertain to the network. However, the semantic layers layout is useful in visualizing the hierarchical nature of this network, integrating TF, non-coding RNA (ncRNA), miRNA and protein-protein interaction data (Fig 4D and Supplementary Video 2). In this figure, network connectivity and hierarchy reflects genomic properties:top level TF-binding correlates with target expression, mid-level contains ‘information flow bottlenecks’ and connections with miRNA and distal regions, revealing ideal drug targets. Such multi-layered heterogenous information integration assists in differentiating intra-level interconnections as well as inter-level edge types and node labels. Note that nodes in each layer are also arranged in 3D using 3D force-directed layout.

**Example 3.** Visualizing the global network of interactions while scaling or coloring a subset of the nodes based on their specific properties can enable hypothesis support. In this example, the visualization helps support the principle that functionally significant and highly conserved genes tend to be more central in physical protein-protein and regulatory networks^52^. Based on this hypothesis, Fig. 4C visualizes a network of tolerance to loss-of-function (LoF) mutations and evolutionary conservation, with nodes for (LoF) tolerant (blue) and essential genes (red) easily distinguishable^52^. Node size is based on the degree centrality of a gene^7^. While essential genes tend to be bigger and central, LoF-tolerant genes are smaller and located in the periphery. Both the 2D snapshot (Figure 4C) and 3D Supplementary Video 1 provide clear visualizations of this complex data that lead to easy interpretation. Note that we have published an iCAVE-generated visualization of a network with similar properties that enabled help support this hypothesis ^53^.

### Graph Clustering To Identify Network Motifs

Clustering is critically important during network exploration, as biomolecules that cluster together tend be functionally related. iCAVE offers the following graph clustering algorithms:

*Edge-Betweenness clustering (EBC)*. The number of shortest paths going through a particular edge is EB. An edge with a high EB value connects multiple communities. At each step, the EBC algorithm removes the edge with the highest EB value until it has optimized a modularity metric on how unlikely the in-cluster degree of a node is in comparison to a random edge. EBC^54^ is an attractive algorithm since it does not require an estimate of the number of clusters *a priori*, unlike a majority of existing graph clustering algorithms.

*Markov clustering (MCL)*^55^ is a scalable and unsupervised algorithm which assumes that the number of intra-cluster connections is large and inter-cluster connections is small. It is based on a bootstrapping procedure that simulates random walks (flow) through the network that expands or contracts in parallel with regional connectivity.

*Modularity clustering (MC)* uses the first eigenvector of the modularity matrix to assign nodes to clusters^56^. While ideal for weighted networks, MC delivers intuitive layouts for networks that do not have weights as well.

### Layout Options for Cluster Visualization

ICAVE can easily visualize the clusters generated by iCAVE or another tool. By default, each cluster is positioned in space with *force-directed layout*^30^, analogous to node positioning. Every cluster is embedded inside a transparent bubble, with members and their connections organized using the hemispherical layout. This arrangement provides a visual aesthetic, and (optional) edge bundling further clarifies the global topology (i.e. thicker bundles for high intra-cluster connectivity). Users can choose alternative layouts for cluster bubble positioning. *Lin-log cluster layout* is a variation of the force-directed model^30^, where highly connected clusters are arranged in closer proximity. *Circos cluster layout* is an innovative algorithm we developed as a 3D adaptation of the popular 2D Circos layout^57^. In this algorithm, we arrange the nodes in 3D space as in hemispherical layout, where the most connected node is located at the center of the hemisphere. We then slice the hemisphere with (pie-like) panels that correspond to separate clusters. Cluster representations can be optimized by variations in node/ slice colorings or edge bundling. **Fig. 6** illustrates different cluster layout options using a metabolite network example.

**Figure 6.**
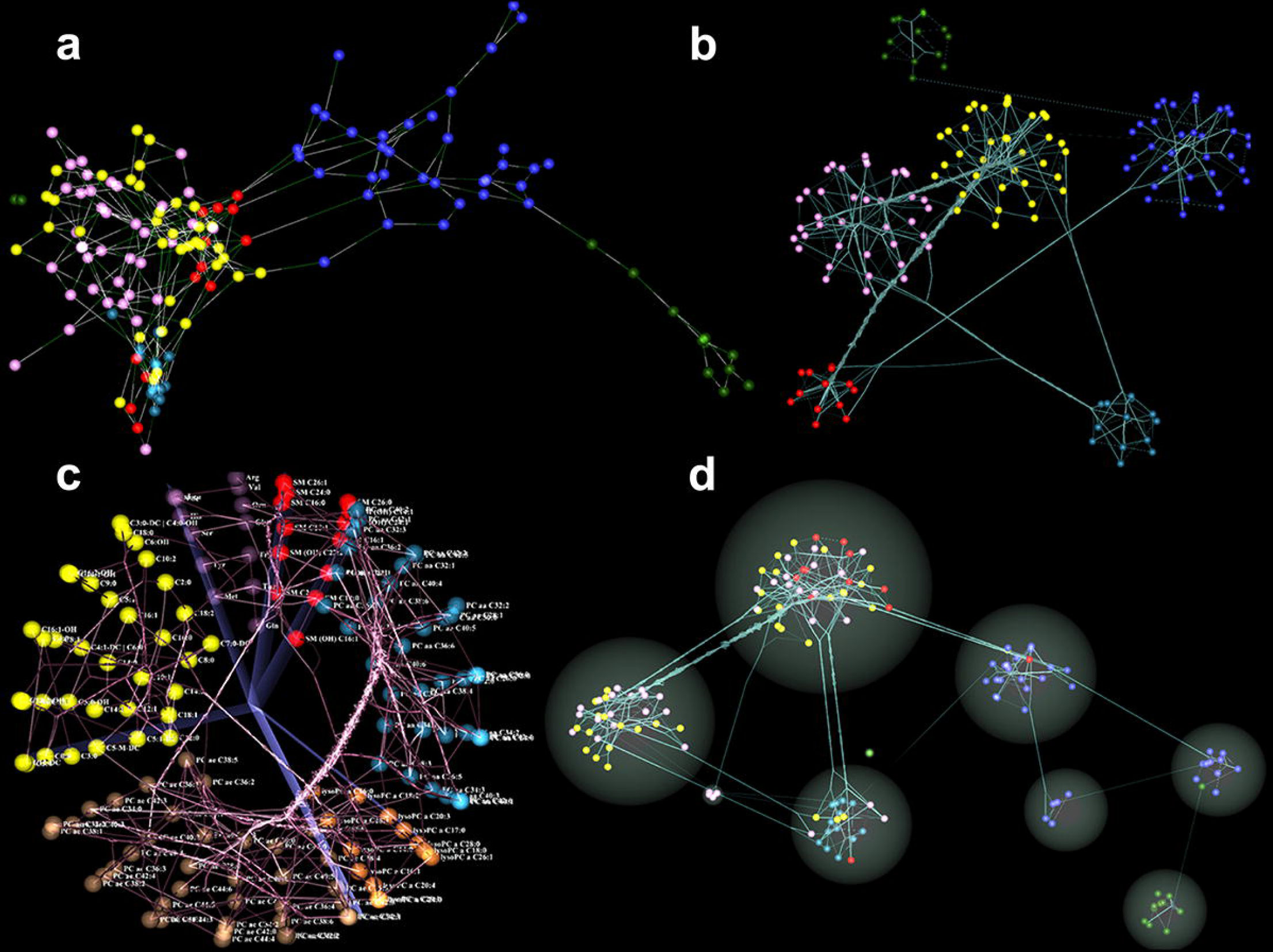
Pathway reconstructed high-throughput metabolomics data with Gaussian Graphical Modeling (GGM)^73^; each sphere color represents a single metabolite class. **a.** The force-directed layout of the weighted network captures the local cluster structures (snapshot). **b.** Snapshot of user-defined metabolite clusters: cluster layout is force-directed while inside each cluster, nodes are ordered in hemispherical layout. Edge bundling represents inter-cluster connectivity strength. **c.** User-defined clusters of the same network in a Circos layout. We provide a movie to better investigate the network in the Supplementary Video 4. **d.** Markov Chain Clustering of the same network based on its connectivity, available as one of the clustering options in iCAVE. Each cluster is represented inside spherical bubble. While topology suggests that most similar metabolites cluster together, this is not always the case, as shown. In all panels, addition of metabolite labels is user-optional.

***User Interface*.** Investigators can easily toggle between alternative layouts of a single graph to emphasize different network aspects. Users without stereo equipment can rotate, zoom or scale the visual to investigate special structures, print 2D snapshots and save movies of a rotating network. Rotation allows multiple views for users without 3D. Exporting and exchanging such movies is very convenient in the YouTube era, enabling easy publication and sharing with collaborators without iCAVE or stereo. Those with a stereo-enabled computer (or a CAVE facility) wear stereoscopic LCD shutter glasses that convey 3D image and allow immersive interaction. In a CAVE, sensors track the user’s eye position and adjust perspective according to user movements. The mouse (or wand) gestures are mapped to logical events that the network layout application handles. Zoom and rotate options activated with a simple mouse (or wand) click help focus on a particular node or edge.

***Shape, color, texture for easier visual data dissemination***. In iCAVE visualizations, 3D spherical glyphs represent nodes. Node color, size and texture optionally encode further statistics (e.g. color for gene induction or repression, size for the magnitude of change in expression, texture to differentiate classes). Edges can be colored, patterned or directed.

***Input/output*.** Input to the iCAVE software is provided as a tab-delimited text file of identifiers and optional information on magnitude of change, edge directionality, edge weights, node/edge colors and patterns. Supplementary Table 2 includes the complete options list. Interaction data are read from an SQLite database. The user can modify the network in real time and store it in DB Browser (which is a light GUI editor for SQLite databases) as a ․db file, so that it can be saved for later access. Output is the layout of the network drawn in VRUI environment, which can be saved as high-resolution 2D image snapshots (.png format) or movies (.gif format).

## Discussion

ICAVE is a freely available open-source biomolecular network visualization tool that leverages advanced (immersive) 3D display technologies and offers several display options integrated with an effective user-interface. It incorporates a number of built-in network layout and graph clustering algorithms to enable automatic generation of 3D visualizations. Based on prior knowledge, input can additionally include (i) 3D node positions; (ii) cluster memberships; or (iii) multi-level hierarchies; or (iv) edge directionality. Utilizing iCAVE, investigators from diverse fields can gain insights from large, heterogeneous datasets, and optimize the quality of their visualizations using different node color, size, transparency options as well as various weight, thickness, transparency and directionality options for the edges. Network topological properties and centrality are also reported. While not extensive, the COMBO database enables disease researchers for quick query of their interactions among genes, drugs and disease phenotypes.

We have designed iCAVE with a modular software structure to create a general and flexible community resource. Users (with some programming experience) can add algorithms for network layout, cluster layout or graph clustering without affecting the core functionality of the code. Users can also add their specific datasets of interest to COMBO. Note that in future iterations of iCAVE, we will implement a Cytoscape^20^ API.

Software, user manual and tutorial are freely available for download to academic users at http://research.mssm.edu/gumuslab/software.html released under the GNU Lesser General Public License.

## Materials and Methods

**Input/Output Formats.** iCAVE supports tabular input formats (.txt, ․csv, or ․tsv). Interactions are either user-defined, or are queries of iCAVE COMBO database. Optional weights are represented with edge color frequency, directed edges with arrows, and node types with node glyph patterns. Input file options are listed in User’s Manual. Networks are saved as static high-resolution (.png) images, or movies of the rotating 3D image (.gif).

**Implementation.** iCAVE uses Virtual Reality User Interface (VRUI), a development toolkit for interactive high performance VR applications^44^, which enables quick and scalable production of completely platform-independent software. iCAVE is thus portable between Linux and Mac system computers (optionally equipped with stereo capabilities) and CAVE facilities.

**Programming Libraries.** Several programming libraries provide intuitive and user-friendly rendering solutions. Vrui library uses a C++ based OpenGL API platform that simplifies handling navigation transformations, light sources, menu creation, and rendering different objects. The SQLite3 software library handles large-scale database parsing. igraph library functions solve some of the programming challenges in generating regular and random graphs, manipulating graphs as well as assigning attributes to nodes and edges. The ANSI C programming language library Argtable enables parsing user-defined 3D graphics options.

**Adding New Algorithms.** Node and edge data are stored in two separate *structure arrays. Node structure* stores its id, name, number of neighbors, color, texture, cluster, size and coordinates. *Edge structure* includes start node id, end node id, weight and color. Storage with structure arrays simplifies the addition of new layout algorithms, because the arrays can be used as inputs. After layout coordinates are calculated, iCAVE utilizes OpenGL API for visualization. New algorithms are added as separate ․cpp files and the corresponding header files are imported to the main program (vrnetview.cpp).

**Label Creation.** Since VRUI offers limited label creation options that render low quality and unreadable text, we developed texture mapping for high quality rendering. Supplementary Figure 1 illustrates VRUI vs. iCAVE labels.

## User interface

Multiple functionalities demonstrate natural modes of interaction for effective analysis. These include activities such as selecting objects and interacting with the image in 3D space. While learning a new user-interface motif has been a traditional weakness of VR environments, more mature and practical technologies are becoming pervasive in consumer markets (e.g. motion sensors in Wii game consoles). These developments inform our user-interface design and provide new users with familiar gestures and interaction motifs.

**Network exploration interface.** Several features enable exploring, interacting and modulating the networks in real-time and saving the result. Interactive menu options are listed in Supplementary Table 2.

**User interface in CAVE environments.** Investigators enter a CAVE environment wearing stereoscopic LCD shutter glasses that convey 3D image. When the user walks around, sensors track movements and the video adjusts accordingly. Multiple users can exist simultaneously in the network and view the visualizations from multiple perspectives by moving in the space, or directly interact with specific biomolecules by clicking on the handheld device to display all its interactions in that network or stored in the database. User can alternatively investigate the network on his own computer.

**Output image generation.** iCAVE assembles image snapshots from several viewpoints into one high-resolution (.png) image (see Supplementary Fig. 2). The desired resolution is user-adjustable via a zoom factor.

## Network Topological Properties

ICAVE automatically calculates the following network properties, rank-orders nodes based on these and represents their distribution both graphically and in tabular form:

**Node degree property** yields hubs. Generally, only a few biomolecules (hubs) have many network interactions^58,59^. Hubs are often central in mediating interactions among the less connected biomolecules ^60,61^.

**Neighborhood connectivity metric** assists in identifying modularity, where small interconnected subgraphs may potentially represent specific enzymes, structures or processes^62,63^ and provide significant insights to perturbed disease mechanisms. For example, the degree of gene coexpression correlates strongly with the complexity of an embedded motif^64^.

**Network average and local clustering coefficients** quantify connectivity of the whole network or a single node. Local clustering coefficient is the ratio between the number of edges that connect the neighbors of a node versus the maximum possible number of edges. The network average clustering coefficient is the average of the local clustering coefficients of all nodes^65^. Only nodes that belong to networks with >3 nodes are considered. The range of coefficient values varies from 0 (no interconnection), to 1 (perfect interconnection).

**Network closeness centrality and node closeness centrality** quantifies the velocity of information flow within a network (the reciprocal sum of the shortest paths from a selected node to all other nodes^66^). Only nodes in subnetworks with >3 nodes are evaluated. When shortest paths are calculated, each edge is scaled with corresponding weight, which can be a floating value. The average of all node closeness centrality values is the network closeness centrality value.

**Network diameter** is the length of shortest path between two farthest nodes. Unconnected nodes are not considered. Irregular networks usually have small diameters, while regular networks have large diameters.

**Betweenness centrality** is a global metric on the importance of a node, which is equal to the number of shortest paths from all vertices to all others that pass through that node, calculating the *load* on a node^67^. Real world scale-free networks usually involve short path lengths across the network, and a few nodes have high betweenness-centrality. *Connector* or *high-traffic* biomolecules that are vulnerable to targeted attacks, usually suggest potential non-hub drug targets^68,69,70^.

**Shared nearest neighbors:** A similarity metric based on the sharing of nearest neighbors between any two nodes. Particularly useful in network topology-based motif, sub-graph or cluster identification.

**Shortest paths:** Quantifies the importance of a node within the network, calculated by the number of shortest paths going through the node. Purely random graphs exhibit a small average shortest path length (~ the logarithm of the number of nodes] along with a small clustering coefficient.

## Layout Algorithms

A graph *G(V ={1,…,n},E)* represents a binary relation *E* over node set *V*. iCAVE both extends classical layouts to 3D and offers novel algorithms. Based on the underlying topology, a user can choose the best layout that helps with data interpretation.

1. **Force-based layout.** The forces acting on each node in classical Fruchterman-Rheingold (FR) algorithm^30^ are:

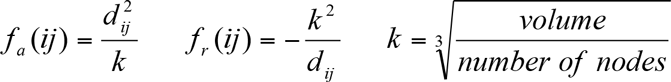

where *f_a_(ij)*and *f_r_(ij)* are attractive and repulsive forces, *d_ij_* is the distance between nodes *i* and *j*, and *k* is a constant corresponding to the equilibrium edge length.
2. **Lin-log layouts.** We used r-PloyLog^41^ energy model to implement the node-repulsion and edge-repulsion LinLog models. For all *r*∈*R* with *r>0*, the node-repulsion energy of a layout *p* is:

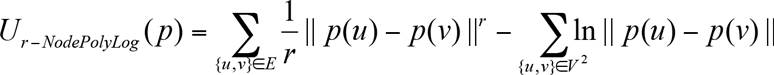

where *p(u)* is the position of node *u*. Edge-repulsion energy is:

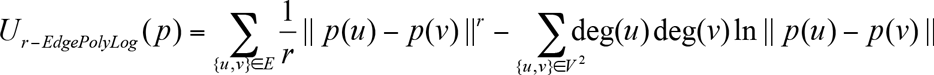

where *deg(u)* is the number of edges incident to node *u*. At *r=3*, the 3-PolyLog reduces to FR and at *r=1* to LinLog model. LinLog models group nodes according to cut density and the normalized cut, therefore the layout leads to graph clustering.
3. **Hemispherical layout.** We place *n* nodes of a graph *G(V={1,‥,n}, E)* equally spaced on a single 3D hemisphere surface, reducing the problem to finding a *hemispherical node ordering*. Coordinates for a node *i є V* are *(x_i_, y_i_, z_i_) є R*, at fixed hemisphere radius R:

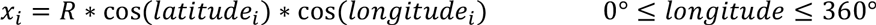

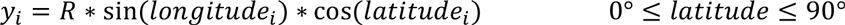

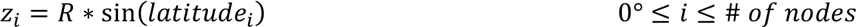

Nodes are sorted and placed based on their degree, with the highest degree node at the hemisphere surface center. Algorithm inputs are the number of nodes, the graph center position and hemisphere radius. Hemisphere radius, node sizes, colors and textures are adjustable.
4. **Semantic levels layout** is ideal for integrative analysis of multiple data resources (e.g. genotype, phenotype, drugs, proteins, metabolites]. Initially, FR algorithm is performed in 2D. Then, multiple equidistant levels (default =7] are created in the z-dimension. Based on network topology, we consecutively assign the nodes to one of the layers. iCAVE user-interface allows the manual manipulation of the number of layers and the distance between them. If layers are not predefined, we suggest experimenting with different options.
5. **Hybrid force directed layout**^42^ Original version of this algorithm is extremely computationally intensive, so we implemented a simplified version, reducing the run time at the expense of visualization quality. Our version has three steps: (i) position nodes randomly; (ii) partition the resulting graph; (iii) apply FR^30^ algorithm separately on each subgraph. The partitioning step splits the graph into two sub-graphs (A and B) of equal sizes. This requires minimizing the cut size, by calculating the second Eigenvector (Fiedler vector) λ of the following:
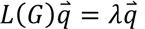 where 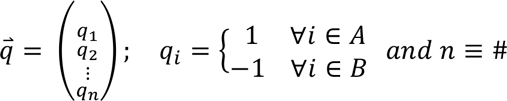 *of nodes* and *L(G)* is the Laplacian of graph *G*. The power-iteration algorithm solves for λ.

**Edge-Betweenness (EB) Clustering Algorithm:** An edge with a high *EB* value potentially connects two or more *communities*. The edge with the highest *EB* value is removed at each step. The number of edges to be removed is user-defined ( with a default of *0.2 times the number of edges)*. Any edge that leads to a single-node cluster is not removed.

**Edge bundling algorithm** is based on application of forces (electrostatic and spring) on an edge subdivided into multiple points. Edge compatibility metrics edge angle, scale (length), position and visual compatibility are multiplied for total compatibility. If two edges are compatible above a threshold, forces are calculated and added to each subdivision, and those subdivisions are bundled together.

## Acknowledgements

This work was supported by the Concern Foundation Conquer Cancer Now Award (to Z.H.G) and the computational resources and staff expertise provided by the Department of Scientific Computing at the Icahn School of Medicine at Mount Sinai, as well as by the PBTECH staff expertise of HRH Prince Alwaleed Bin Talal Bin Abdulaziz Alsaud Institute for Computational Biomedicine and the computational resources of the Coffrin Center for Biomedical Information at Weill Cornell Medical College of Cornell University (to V.L. and Z.H.G). We wish to thank Dr. Jian Sun and Jason Banfelder for programming assistance and constructive comments. To ensure that the proposed visualization platform meets the needs of translational and basic science researchers, we relied heavily on the advice, feedback and discussions from representative end-users including Mark Gerstein, Alex Lash, Michael LeVine, Chris Sander, Eric Schadt and Karsten Suhre.

## Author Contributions

Z.H.G. conceived the study, designed all the analyses and oversaw all aspects of the project; V. L. wrote most of the code with S. K., A. G. and M. W. and V. L. generated the figures. Z.H.G. wrote the manuscript with contributions from all authors.

